# Widespread Horizontal Gene Transfer Among Animal Viruses

**DOI:** 10.1101/2024.03.25.586562

**Authors:** Christopher B. Buck, Nicole Welch, Anna K. Belford, Arvind Varsani, Diana V. Pastrana, Michael J. Tisza, Gabriel J. Starrett

**Affiliations:** National Cancer Institute, Bethesda, MD, USA; Arizona State University, Tempe, AZ, USA; University of Cape Town, South Africa

**Keywords:** coronavirus, nidovirus, maldvirus, meldvirus, spiretrovirus, tailtomavirus, adabscovirus, helpol, rep, replicase, cap, capsid, virion, hexon, penton, macc, cah, adenain, bidnavirus, adenovirus, adomavirus, papillomavirus, parvovirus, polyomavirus, papyomavirus, poxvirus, herpesvirus, iridovirus, megavirus, giant virus, mimivirus, anellovirus

## Abstract

The initial objective of this study was to shed light on the evolution of small DNA tumor viruses by analyzing *de novo* assemblies of publicly available deep sequencing datasets. The survey generated a searchable database of contig snapshots representing more than 100,000 Sequence Read Archive records. Using modern structure-aware search tools, we iteratively broadened the search to include an increasingly wide range of other virus families. The analysis revealed a surprisingly diverse range of chimeras involving different virus groups. In some instances, genes resembling known DNA-replication modules or known virion protein operons were paired with unrecognizable sequences that structural predictions suggest may represent previously unknown replicases and novel virion architectures. Discrete clades of an emerging group called adintoviruses were discovered in datasets representing humans and other primates. As a proof of concept, we show that the contig database is also useful for discovering RNA viruses and candidate archaeal phages. The ancillary searches revealed additional examples of chimerization between different virus groups. The observations support a gene-centric taxonomic framework that should be useful for future virus-hunting efforts.

## Introduction

It is well established that different virus families can exchange genes via recombination (Bennett et al., 2010; Koonin et al., 2015). Examples of horizontal genetic modularity can be found among small DNA tumor viruses, a polyphyletic group traditionally encompassing adenoviruses, papillomaviruses, parvoviruses, and polyomaviruses. An emerging group of fish-tropic viruses called adomaviruses unite <u>ad</u>enovirus-like virion proteins with poly<u>oma</u>virus- or papill<u>oma</u>virus-like replicative DNA helicases (Dill et al., 2018; Mizutani et al., 2011; Welch et al., 2020). Another emerging group called adintoviruses unite <u>ad</u>enovirus-like replicase and virion proteins with retrovirus-like <u>int</u>egrase genes (Starrett et al., 2021).

The widespread horizontal transfer of genes among virus groups differs from the typical pattern observed for eukaryotes, which evolve primarily through vertical inheritance. The crisscrossing evolutionary paths of viral gene modules present a major challenge for traditional Linnaean whole-organism taxonomic approaches. For example, bandicoot-tropic papyomaviruses (which unite <u>pa</u>pillomavirus virion protein genes with a <u>p</u>ol<u>yom</u>avirus replicase gene (Bennett et al., 2010; Woolford et al., 2007)) are <u>unclassifiable</u> under current taxonomic approaches.

The starting goal of this study was to detect divergent small DNA tumor virus sequences in datasets available through the National Center for Biotechnology Information (NCBI) Sequence Read Archive (SRA). Iterative rounds of searches for homologs of individual genes of interest revealed an increasingly diverse range of sequences representing additional virus groups. Because many of the contigs identified in the survey appear to represent chimeras, the annotation process required us to develop a gene-centric taxonomic framework that we hope can facilitate future virus discovery and classification efforts.

## Methods

### Datasets

Short-read deep sequencing datasets were selected using web-based <u>SRA</u> <u>searches</u>. In some cases, entire BioProjects of interest were subjected to *de novo* assembly. In other cases, un-assembled read sets were scanned for small DNA tumor virus hallmark protein sequences of interest using <u>Diamond</u> 2.0 (Buchfink et al., 2021) with settings --block-size 7 --index-chunks 1 --evalue 0.00001 --outfmt 6 qseqid sseqid evalue qseq. SRA datasets with relatively large output .tsv files (generally a >3 kb file size cutoff) were selected for *de novo* assembly.

Several dozen new viral genome sequences were detected through re-analyses of our group’s previously published rolling circle-amplified (RCA) metagenomic surveys (Nguyen et al., 2017; Pastrana et al., 2018; Tisza et al., 2020). Maps of contigs in this local direct-sequencing category have file names ending with “Run#s#” (File1_NamingKey.xlsx and File2_Maps.zip). A subset of genomes were found in datasets offered by the <u>Joint Genome Institute</u>, <u>Wild Biotech</u> (Levin et al., 2021), or in tBLASTn searches of NCBI WGS and TSA databases. GenBank does not allow third party annotated submissions derived from non-SRA databases, so names of maps in this category begin with the lowercase letter u (unsubmitted).

### Sequence processing and de novo assembly

Sequences were initially downloaded using the fastq-dump module of <u>SRA Toolkit</u> 2.9.6, with settings -X 50000000 -I --split-files --gzip. More recent assembly efforts used the fasterq-dump module of SRA Toolkit 3.0.3 to download reads and <u>Seqtk</u> was used to sample 50 million read pairs with the setting -s100. Reads were quality-trimmed using <u>fastp</u> (Chen et al., 2018). <u>Megahit</u> 1.2.9 was used to assemble the trimmed reads with setting --min-contig-len 1500 (Li et al., 2015). A <u>python script</u> was developed to rename contig fasta headers based on the accession number of the parent SRA dataset.

### Virus detection and gene annotation

In most cases, *de novo*-assembled contigs were imported into <u>CLC Genomics Workbench</u> 22 and converted into BLAST databases. The contigs were searched using tBLASTn against an iteratively refined collection of protein baits representing hallmark virus genes of interest. Assemblies were also subjected to Diamond screens to detect specific protein sequences of particular interest. Contigs of interest were examined using <u>BLASTx</u> searches against GenBank nr_clustered taxid:10239 (viruses) or were detected using Cenote Taker 2 (Tisza et al., 2021). Contigs were manually validated and extended using the Map Reads to Reference function of CLC Genomics Workbench 22. <u>MacVector</u> 18 was used to compile annotations and to display maps. Predicted protein sequences are compiled in File3_AllProteins.fasta and File4_AllProteinsE5.cys. The latter file can be viewed using <u>Cytoscape</u> software (Shannon et al., 2003).

Contigs inferred to represent viral sequences endogenized into host genomes, contigs with insufficient read coverage to assemble a complete viral genome, sequences with close homologs already represented in GenBank nr, and virus groups outside the scope of the current survey were auto-annotated using <u>Cenote Taker 3</u> with settings -db virion rdrp dnarep -hh hhsearch and --caller phanotate. Maps for contigs in this “unsubmitted” category begin with the lowercase letter “u” (File 1_NamingKey.xlsx and File2_Maps.zip). Manually re-annotated maps of previously published reference viral genomes are flagged with the initial term “ref.”

Translated ORFs with clear hits (E values < E-05) in NCBI DELTA-BLAST searches were assigned a common gene/protein name. An example is the papillomavirus replicative helicase, homologs of which have clear hits that universally share the name “E1.” When new homologs were found in different virus families (e.g., papillomavirus E1 homologs found in cressviruses and adomaviruses) the gene was assigned a descriptive hybrid name (e.g., RepE1). For genes with multiple existing names, a descriptive name was developed and appended with an element of a more common gene name. As a specific example, an adenovirus oncogene represented in the literature by various combinations of the terms early E4, protein 6, control protein, 15.9 kDa, 16 kDa, 19.8 kDa, 32 kDa, 33.2 kDa, 34 kDa, 34.1 kDa, 34.6 kDa, 34.7 kDa, 34K, 34K-2, E4.2, E4-1, E4-3, E4-6, BAdVBgp27, ORFD, ORFE, ORF3, ORF5, ORF6, ORF6/7, ORF26, 245R, 253R, hypothetical protein, or E4orf6 was assigned the name “Oncorf6.”

Candidate protein sequences without clear hits in BLAST or Cenote Taker 2 analyses were subjected to <u>HHpred</u> analysis using databases PDB_mmCIF70, Pfam-A, UniProt-SwissProt-viral70, and NCBI Conserved Domains (Zimmermann et al., 2018). Some sequences without clear hits in HHpred searches were subjected to <u>Phyre^2^</u> searches or <u>Dali</u> searches using AlphaFold2 or <u>RoseTTAfold</u> structure predictions (Baek et al., 2021; Holm et al., 2023; Jumper et al., 2021; Kelley et al., 2015). Predicted protein sequences were also scanned for <u>short linear motifs</u> of interest (Kumar et al., 2022).

Sequences with no clear hits in any of the above methods were assigned a distinctive name. When choosing gene names, we generally sought to assign each protein cluster in all-against-all network analyses (File5_PolishedE5.cys, File6_PolishedE10.cys) a distinctive root name consisting of a single pronounceable term starting with a capital letter.

Some gene categories, such as the Cah and Macc classes proposed in the current report, were named based on generic predicted folds, such as alpha helical coiled coil domains. The Oncoid class is defined solely by the presence of short linear motifs of interest. In many cases, unidentifiable proteins were assigned an entirely arbitrary neologism loosely reflecting an HHpred hit (a convention that was used even in cases where all the HHpred hits were very weak). The working hypothesis behind this strategy is that homologs of the hypothetical protein might share some of the same weak HHpred hits, facilitating future recognition of additional homologs. Neologisms are not meant to imply claims about protein structure or function.

Protein sequence clusters were analyzed using <u>EMBOSS getorf</u>, <u>EFI-EST</u> (Oberg et al., 2023; Zallot et al., 2019), and Cytoscape. Phylogenetic trees were inferred with PhyML using <u>Phylogeny.fr</u> One Click mode (Dereeper et al., 2010; Dereeper et al., 2008). Trees were viewed using <u>FigTree software</u>. <u>KnotInFrame</u> (Janssen & Giegerich, 2014; Theis et al., 2008) was used to predict programmed −1 ribosomal frameshift slippery sequences.

### Virus names

This study proposes common names for viruses. Virus category names should be viewed as convenient handles that do not necessarily indicate universal traits of the group. As an example, papillomaviruses were originally named for the ability of the founding members of the group to cause papillomas (skin warts), and the name has been maintained even though subsequent study revealed that most members of the group rarely cause noticeable skin lesions. When choosing names, we have sought to comply with Section 2.1 of the <u>International Code of Virus Classification and</u> <u>Nomenclature</u>: “(i) to aim for stability; (ii) to avoid or reject the use of names which might cause error or confusion; (iii) to avoid the unnecessary creation of names.”

A general philosophy of the current report is that “lumping” virus groups is generally more useful than “splitting.” We broadly define adomaviruses as predicted circular contigs 9-20 kilobasepairs (kb) in length that encode a superfamily 3 helicase, a candidate Penton, and a candidate Cah-class gene (see legend of Figure 1 for a key to major gene categories). Parvoviruses are broadly defined as contigs that encode a parvovirus-like NS1 replicase and one of several broad classes of parvovirus-like virion proteins (VP). The “lumped” definition of parvoviruses encompasses species with proposed Cah domains, species that appear to have circular genomes, candidate segmented species, and species with auxiliary CressRep, PolB, or HelPol replicases. The standard polyomavirus definition has been expanded to include species with proposed Cah domains and species that encode inferred early and late genes on the same strand.

**Figure 1:**
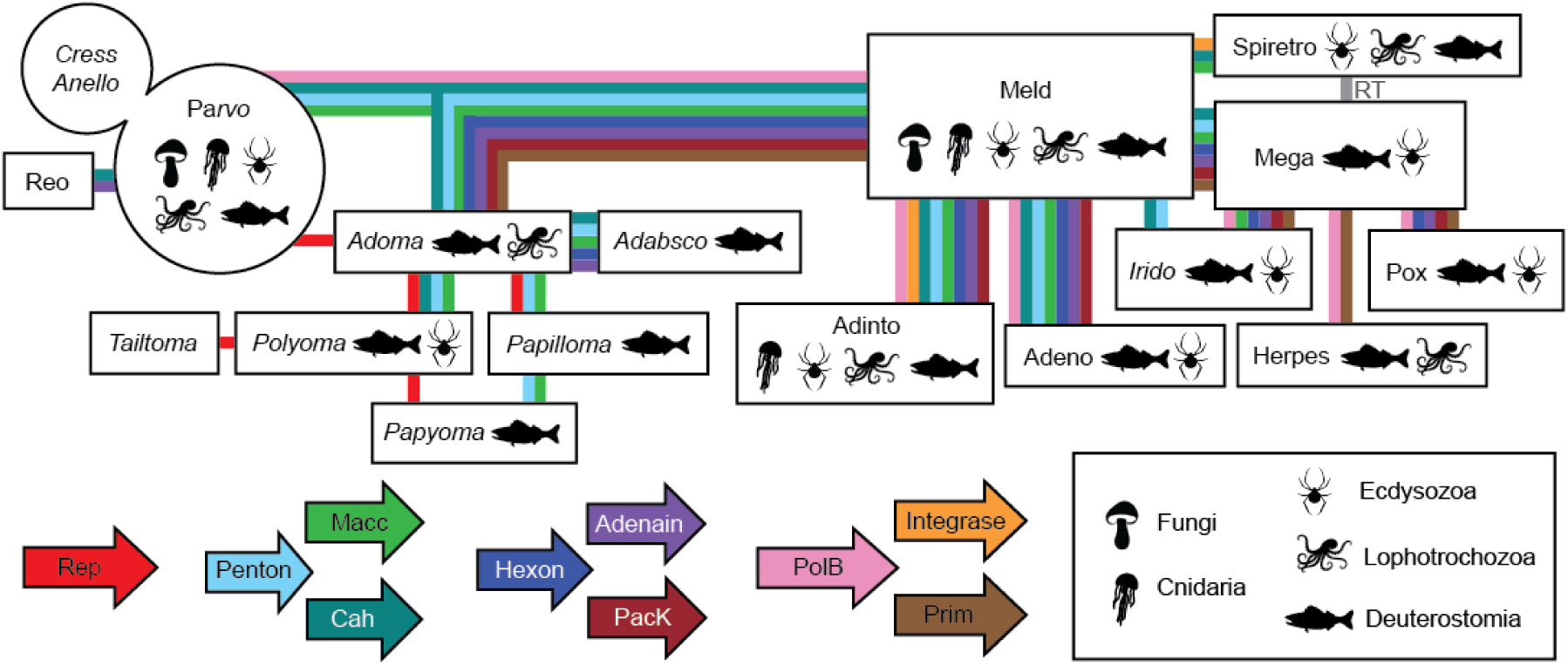
A hypothetical gene flow framework representing the observed distribution of hallmark small DNA tumor virus-related genes observed in opisthokont datasets. Colored lines indicate at least one inferred transfer event for the indicated gene involving at least one member of each virus group. The line does not imply that all members of the group received or maintained the indicated gene. In some cases, the line may represent multiple inferred horizontal gene transfer events occurring at different times. Italics indicate viruses predicted to have circular genomes. **Rep**: superfamily 3 ATP-dependent <u>rep</u>licative DNA helicase with N-terminal nicking endonuclease-like domain. Familiar parvovirus, polyomavirus, and papillomavirus Rep genes are known as NS1, LT, and E1, respectively. **Penton**: pentameric single β- jellyroll capsid protein. Familiar parvovirus, polyomavirus, and papillomavirus genes of this class are known as VP, VP1, and L1, respectively. We speculate that a subset of parvovirus and <u>c</u>ircular <u>R</u>ep-<u>e</u>ncoding <u>s</u>ingle-<u>s</u>tranded DNA virus (cressvirus) capsid genes may represent captured <u>m</u>idsize <u>e</u>ukaryotic linear <u>D</u>NA virus (meldvirus) Penton homologs. **Macc**: hypothetical <u>m</u>embrane-<u>a</u>ctive <u>c</u>apsid <u>c</u>ore protein. Some Macc classes have predicted holin-like hydrophobic domains. Others are predicted to have lipase activity. Adenovirus proteins VI and X are canonical examples of this category. **Cah**: hypothetical <u>c</u>apsid-surface protein with predicted <u>a</u>lpha <u>h</u>elical (particularly coiled-coil) character. Adenovirus protein IX is a canonical example of this class. A typical hit for candidate Cah proteins in HHpred searches is the reovirus outer capsid protein sigma-1. The adomavirus gene initially designated LO4 encodes a protein that qualifies as a Cah. **Hexon**: double β-jellyroll major capsid protein that trimerizes to form virion facets. **Adenain**: papain-like cysteine protease involved in virion maturation. **PacK**: Fts<u>K</u>/HerA-class P-loop ATPase proteins thought to facilitate viral genome <u>pac</u>kaging. **PolB**: viral type B DNA polymerase. **Integrase**: retrovirus-like (rve) integrase. **Prim**: homolog of archaeal-eukaryotic <u>prim</u>ase small catalytic subunits. **RT**: reverse transcriptase.

Where possible, the first term of the suggested common name represents an existing roughly family-level group to which the virus appears to belong. In cases where the contig unites elements of different virus families, we apply the “ad-oma” convention, in which the first half of the name represents the category of major virion proteins and the second half of the name represents the replicase category.

The second word in the suggested common name for each virus species represents the animal genus (or type of environmental sample) indicated in SRA metadata, together with the length (in basepairs) of the initially detected contig. Different variants of the same virus species (or inferred segments of a single virus species) are assigned a shared identifier number based on the length of the first observed variant or segment. In some cases (e.g., the human adenovirus type 5 sequence integrated into HEK 293 cells) the identifier number represents a previously identified type number (Adenovirus homo5).

We use isolate designations to convey ancillary information, such as the common name of unfamiliar host animal genera, the specific tissue sampled, or information about virus subcategories. For example, a virus we previously named “Tilapia adomavirus 2 isolate 4096” (<u>BK010892</u>) would instead be named “Adomavirus oreochromis4096 isolate tilapiaAdPa” - where the term “AdPa” indicates <u>ad</u>enovirus-class virion protein genes united with a <u>pa</u>pillomavirus E1 replicase. Throughout the manuscript, we apply a name abbreviation convention consisting of the virus category name plus the identifier number (e.g., Adoma4096).

Final maps were exported in GenBank format and converted into five-column feature tables using a <u>conversion tool</u> developed by Ramsey and colleagues (Ramsey et al., 2020). Fasta header and comment information were compiled in Excel (see “Submission” tab of File1_NamingKey.xlsx).

### Data availability

Accession numbers for viral genomes deposited into GenBank in association with this work are listed in File1_NamingKey.xlsx (tab BasicInfo). GenBank-formatted text maps are provided in File2_Maps.zip. A Diamond-searchable version of the ∼4 TB contig library assembled for this study will be hosted by the National Cancer Institute’s <u>Cancer Data Service</u>. The contig library is available as a free <u>download</u> via Google Drive. The library will also be distributed upon request via Globus or Aspera.

## Results

### Complex gene flow among DNA viruses

Because many genes discovered in this survey harbored little or no amino acid sequence similarity to known proteins, it was necessary to rely on HHpred analyses as well as Dali analyses of protein structures predicted by AlphaFold2 or RoseTTAFold (File7_PDB.zip and <u>File8_Dali.zip</u>). In many cases, gene neologisms or gene names flagged with the suffix “-oid” represent rough guesses about the possible nature of the encoded protein. Proposed names should not be construed as claims about the actual structures, functions, or evolutionary relationships of predicted genes. We anticipate that some suggested gene names may eventually need to be updated when more information about their biological functions and inferred evolutionary relationships becomes available. As a guide to gene- and virus-naming conventions, we developed a “subway map” representing hypothetical gene flows between small DNA tumor viruses and other virus groups (Figure 1).

The Results section highlights salient examples of proposed annotations from a subset of the 618 GenBank submissions associated with this study (File1_NamingKey.xlsx and File2_Maps.zip). A summary of the gene content for submitted viruses can be viewed interactively as <u>Cytoscape</u> all-against-all network analyses (File5_PolishedE5.cys and File6_PolishedE10.cys).

### Cressviruses

It has previously been proposed that the nicking endonuclease/superfamily 3 replicative DNA helicase (Rep) genes of papillomaviruses, parvoviruses, and polyomaviruses evolved from the Rep genes of a diverse group of small eukaryotic viruses called cressviruses (phylum *Cressdnaviricota*)(Kazlauskas et al., 2019). In support of this concept, the survey detected several candidate cressviruses encoding Rep protein sequences similar (BLASTp E-values 1E-06 to 1E-12) to the <u>E1 rep</u>licase proteins found in papillomaviruses (Figure 2). A RepE1-bearing cressvirus sequence, Cressvirus paraphelliactis4479 (<u>Cress4479</u>), was assembled from a <u>dataset</u> representing muscle tissue from a *Paraphelliactis xishaensis* sea anemone (Feng et al., 2021). The contig also encodes a Hepecap protein with a predicted fold distantly similar to the <u>cap</u>sid proteins of <u>hep</u>atitis <u>E</u> virus and the ssRNA phage MS2 (HHpred probability=67%, AlphaFold-Dali Z-score=6). Other contigs assembled from the same anemone dataset unite fragments of Cress4479 with typical cnidarian genomic sequences, including high-abundance retrotransposon-like sequences (uCress2996, 5451, 6486, and 11852). Another possible case of RepE1 cressvirus endogenization was observed in a <u>dataset</u> representing an *Anemonia viridis* sea anemone (uCress1997). The appearance of high-copy, highly rearranged integration events is reminiscent of patterns observed in tumors caused by papillomaviruses and polyomaviruses (Akagi et al., 2023; Starrett et al., 2020).

**Figure 2:**
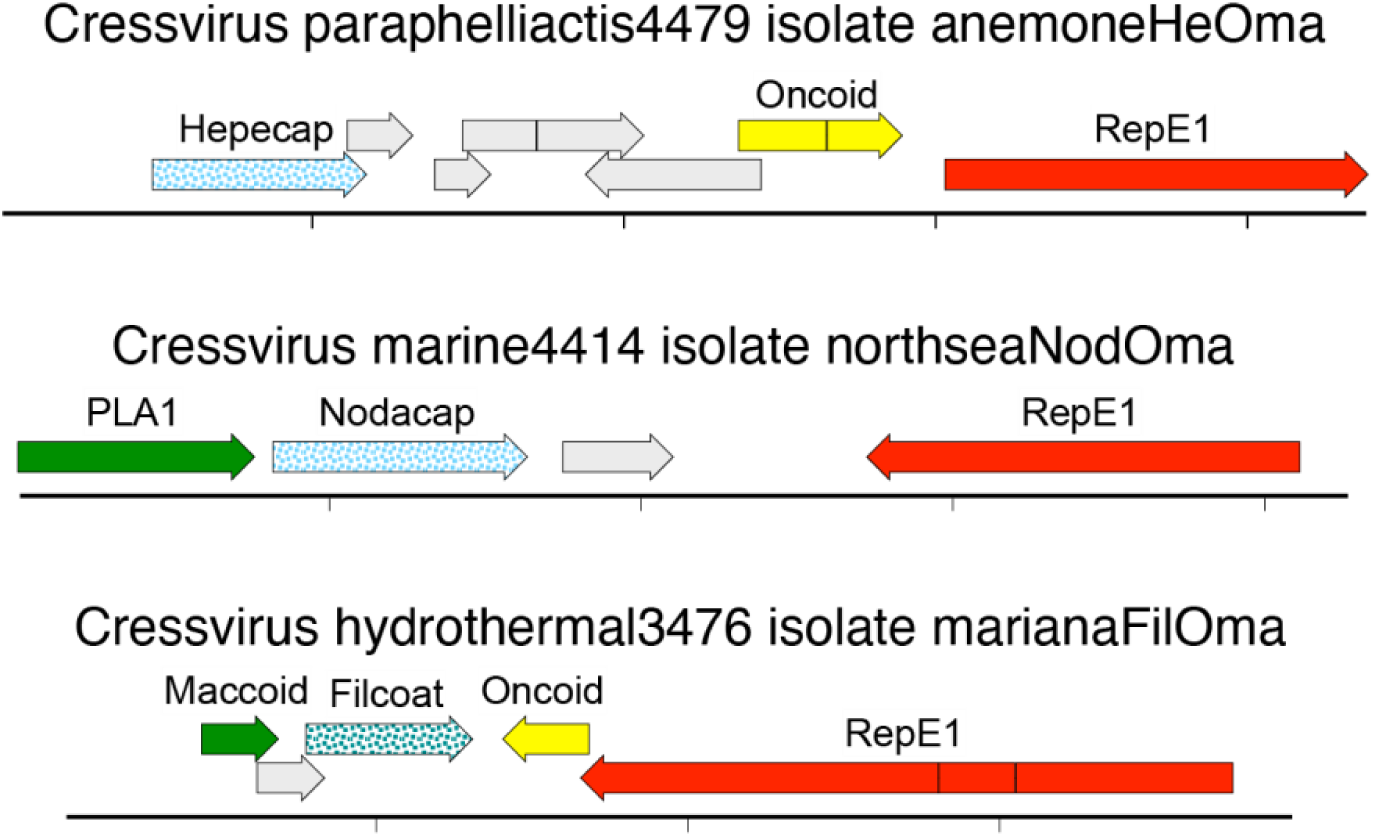
Candidate cressviruses encoding papillomavirus-like replicase proteins (RepE1) paired with different classes of inferred capsid proteins. Putatively circular genomes are linearized for display purposes. Hatch marks indicate one kilobase.

An apparently circular RepE1-bearing sequence from a <u>seawater sample</u>, <u>Cress4414</u>, encodes a candidate capsid protein, Nodacap, with a predicted fold similar to <u>noda</u>virus <u>cap</u>sid proteins (Z=18)(Figure 2). A similar contig, <u>Cress4399</u>, was observed in a *Delisea pulchra* (red algae) <u>dataset</u>. We previously reported (Tisza et al., 2020) a <u>red</u> <u>snapper-associated cressvirus</u> sequence with a similar Nodacap late region, but that sequence carries a traditional cressvirus Rep in place of the RepE1 observed in Cress4414. Another RepE1-bearing cressvirus sequence from a deep-sea <u>hydrothermal</u> <u>vent sample</u>, <u>Cress3476</u>, encodes a protein with a predicted alpha helix-rich fold resembling the <u>fil</u>amentous major <u>coat</u> proteins of archaeal viruses (Filcoat, Z=8).

### Parvoviruses and candidate midsize archaeal linear DNA viruses

A group of parvovirus-like sequences traditionally referred to as bidnaviruses are defined as uniting a parvovirus superfamily 3 replicative DNA helicase (NS1), a parvovirus single-jellyroll capsid protein (VP), and an Alpha-class adintovirus type B DNA polymerase (PolBa) (Starrett et al., 2021). Although some members of the bidnavirus group appear to be monopartite, others have up to twelve genome segments (Wallace et al., 2021). Segmented species typically encode a <u>Cah</u>-like protein similar to cypo reovirus VP2 proteins fused to a pap<u>ain</u>-like protease domain (Cahain). Cahain-bearing genome segments often also encode genes resembling the thymidine <u>k</u>inase (Tkoid) or d<u>UT</u>Pase (Utoid) enzymes of larger DNA viruses. In the current survey, bidna-class <u>parvovirus segments</u> were detected in a <u>dataset</u> representing *Lactarius* fungi. Some contigs assembled from this dataset (uParvo5312e1, 5312e2) appear to represent integrated bidna-class parvovirus sequences adjacent to fungal retroelements or other inferred fungal genomic sequences.

A group of relatively large parvoviruses (genomes >5 kb) encode candidate Cah proteins. Surprisingly, it appears that most members of the Cah-bearing parvovirus group may have circular genomes rather than the expected linear genome flanked by inverted terminal repeats (a hallmark of classic parvoviruses). A different parvovirus group, represented by <u>Parvo3071</u>, encodes an auxiliary CressRep alongside the expected parvovirus-class NS1. These observations blur the lines between the parvoviruses and cressviruses.

The current survey revealed a group of parvoviruses with an exotic DNA polymerase showing remote similarity to cellular PolBc and Pol-Delta proteins (BLASTp 6E-12). A representative example of the group is shown in Figure 3. A Diamond screen of the contig library using the exotic DNA polymerase sequence as bait recovered contigs with mixed similarities to adintoviruses and other <u>m</u>idsize <u>e</u>ukaryotic linear <u>D</u>NA viruses (<u>Meld24113</u>, <u>16152</u>, <u>25100</u>, <u>10171</u>, <u>10183</u>, <u>29240</u>)(Starrett et al., 2021). Sequences distantly similar (BLASTp 2E-06) to the exotic DNA polymerase have recently been reported in viruses thought to infect methanotrophic archaea that inhabit deep-sea hydrothermal vents (e.g., <u>PBV266</u>)(Laso-Pérez et al., 2023). We suggest the gene name HelPol for the exotic DNA polymerase, after the Norse goddess of the underworld.

**Figure 3:**
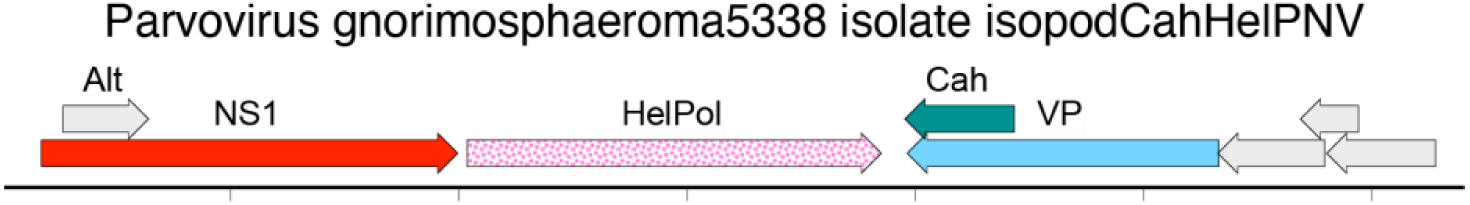
A parvovirus-like sequence encoding an exotic HelPol DNA polymerase.

A dozen highly divergent HelPol-bearing contigs were detected in environmental datasets, including hydrothermal vent samples. We hypothesize that these sequences represent <u>m</u>idsize <u>a</u>rchaeal linear <u>D</u>NA viruses (maldviruses). Analysis of the candidate HelPol maldvirus contigs suggests a range of possible virion architectures, including inferred examples of the familiar adenovirus-like Hexon-Penton organization as well as various combinations of other candidate virion proteins provisionally designated Prop, Protohexon, Recex, and Tailon. These observations suggest a high degree of genomic plasticity among HelPol-bearing archaeal viruses.

### Adomaviruses and tailtomaviruses

The main structural building block of a wide range of viral capsids is a common fold known as an 8-stranded β-jellyroll (Krupovic & Koonin, 2017). The facets of the icosahedral adenovirus virion are formed by trimers of a tandem double-jellyroll protein called Hexon, while the vertices are formed by pentamers of a single-jellyroll protein called Penton. Adenovirus protein IX forms a triskelion and coiled-coil “hairnet” that stabilizes the valleys between Hexon jellyrolls (Gallardo et al., 2021). Core protein VI is thought to emerge during the infectious entry process to play a role in destabilizing host membranes. A papain-like cysteine protease known as Adenain cleaves various capsid proteins during the virion maturation process. Previously reported adomaviruses appear to encode homologs of each of these major adenovirus virion proteins (Welch et al., 2020).

The current data mining effort cataloged more than two dozen complete or nearly complete adomavirus genomes. The group encompasses a surprising degree of genomic diversity, with several classes of inferred replicases and several classes of inferred virion proteins. The adomavirus early (replicase) and late (virion protein) gene modules exist in a range of combinations.

Previously described adomaviruses unite adenovirus-like virion proteins with either a polyomavirus-like replicase (RepLT) or a papillomavirus-like replicase (RepE1)(Figure 4). The survey detecta a third class of Rep genes that resemble the replicases of previously reported cressviruses (CressRep). The isolate names of the three classes of adomavirus replicase are appended with -Py (RepLT), -Pa (RepE1), or -Ress (CressRep).

**Figure 4:**
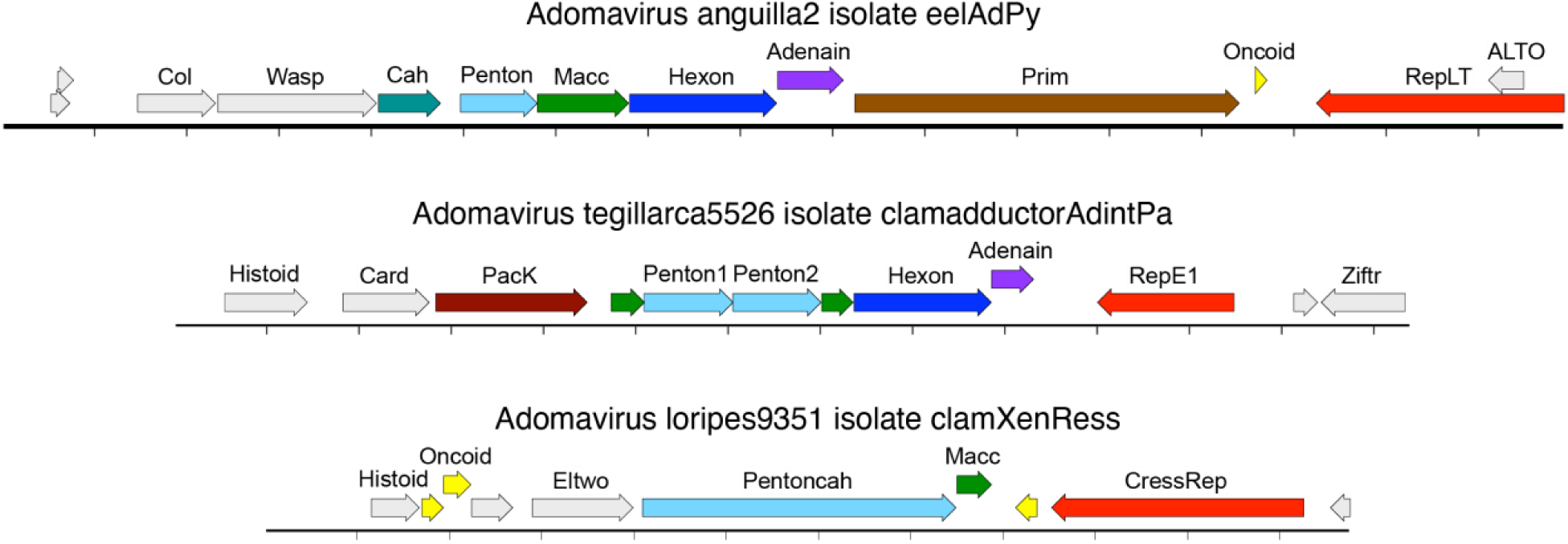
Examples of adomaviruses with three different classes of replicase and three different classes of inferred virion proteins. The circular genome is linearized for display purposes. Hatch marks indicate one kilobase.

Several adomavirus species encode an inferred late region category we initially designated “Xeno” (Welch et al., 2020), denoting of the absence of discernible sequence similarity to any known proteins. Using more recently developed AlphaFold2 and RoseTTAFold structural prediction algorithms, we were able to develop the hypothesis that the Xeno-class late region encodes a highly derived <u>Penton</u> homolog fused to an elaborate candidate <u>Cah</u> homolog (Pentoncah, Figure 4). Xeno-class adomaviruses typically also encode a gene we designate Histoid, denoting its remote similarity to various <u>histo</u>ne proteins (HHpred >95%).

An adomavirus sequence found in a *Tegillarca* clam <u>dataset</u> pairs RepE1 and Histoid genes with sequences similar to adintovirus virion proteins (BLASTp 5E-29)(Figure 4). The early region of <u>Adomavirus tegillarca5526</u> encodes a sequence with remote similarity to CTCF and other <u>zi</u>nc finger transcriptional regulators (Ziftr, HHpred 80%). Ziftr genes are also observed in some adintoviruses (e.g., <u>Adinto12738</u>).

Although the *Tegillarca* clam AdintPa adomavirus appears to encode a homolog of the Fts<u>K</u>/HerA-like ATPase that facilitates the <u>pac</u>kaging of adenovirus genomes, PacK homologs were not detected in other adomaviruses. However, some adomaviruses appear to encode a set of short ORFs upstream of the virion protein genes that could collectively encode protein sequences distantly similar to PacK. In particular, an adomavirus detected in a <u>dataset</u> for a saliva sample from a South American village dog (<u>Adoma16400</u>) encodes an ATP-binding P-loop motif (a hallmark of PacK proteins). We hypothesize that PacK may have been an ancestral adomavirus feature but the gene has been partially or entirely lost in nearly all observed examples.

A seemingly chimeric circular genome detected in a marine <u>dataset</u> pairs a RepLT gene with an inferred late region encoding a protein with a predicted central beta helical domain reminiscent of the adenovirus <u>tail</u>spike-like pr<u>o</u>tei<u>n</u>s E1B-55k and LH3 (Tailon, Z=12)(Figure 5)(Gallardo et al., 2021). In addition to the predicted Tailon protein, the virus also encodes a protein with a predicted <u>ten</u>-stranded β-jellyroll fold similar to pentameric turret pr<u>o</u>tei<u>n</u>s and hexameric single-jellyroll VP4 capsomers of archaeal phages (Tenton, Z=5)(Hartman et al., 2019; Wang et al., 2019). <u>Tailtomavirus</u> <u>marine3478</u> also encodes a Groeloid protein with a predicted fold similar to <u>GroEL</u> and other ring-shaped chaperones (Z=6), as well as a candidate <u>histo</u>ne homolog (Histoid, Z=12), a protein with remote similarity to Z3-type <u>APOB</u>EC deaminases (Apoboid, Z=4), and a candidate oncogene with remote C-terminal similarity to the p53-downmodulating adenovirus <u>onco</u>protein known as E4<u>orf6</u> (Oncoidorf6, HHpred 80%).

**Figure 5:**
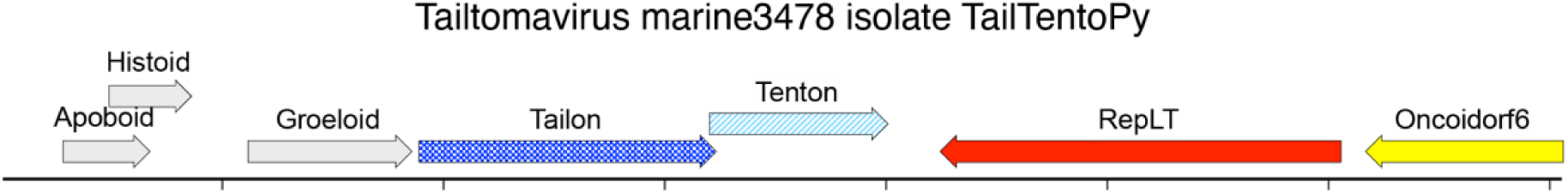
Tailtomavirus detected in a sample of seawater from off the coast of Tierra del Fuego. Dali analysis of AlphaFold-predicted structures suggests tailspike-like and single-jellyroll-like virion proteins unified with a polyomavirus-like replicase protein. The inferred circular genome is linearized for display purposes. Hatch marks indicate one kilobase.

### Anelloviruses and adabscoviruses

Anelloviruses are a group of small single-stranded circular DNA viruses with a single-jellyroll major capsid protein that resembles the major capsid proteins of picornaviruses and avian cressviruses, such as beak and feather disease circovirus (Butkovic et al., 2023; Liou et al., 2022). A puzzling aspect of anelloviruses is that they do not appear to encode any known class of replicase.

Three nearly identical inferred circular genomes were found in <u>three separate</u> <u>bioprojects</u>representing *Betta* fighting fish. The genomes encode <u>ad</u>omavirus-class virion proteins (BLASTp 1E-68) but do not encode a discernible replicase gene (Figure 6). A predicted 70 kDa protein, assigned the arbitrary neologism <u>Absco</u>ndo, shows a ∼40 amino acid C-terminal patch of remote similarity (HHpred 44%) to fowl adenovirus <u>ORF12</u>. Although ORF12 appears to represent a captured parvovirus NS1 Rep gene, the Abscondo sequence lacks the hallmark HUH and P-loop motifs found in known parvovirus replicase proteins. The Abscondo ORF encodes a +1-overprinted leucine-rich sequence reminiscent of polyomavirus Middle T and ALTO genes (Carter et al., 2013). The candidate overprinted gene is assigned the name Arnao, denoting remote similarity to the <u>ar</u>ginine-rich <u>N</u>-terminal domain of <u>a</u>nellovirus <u>O</u>RF1 capsid proteins, gyrovirus VP1 capsid proteins, and beak and feather disease circovirus Cap proteins.

**Figure 6:**
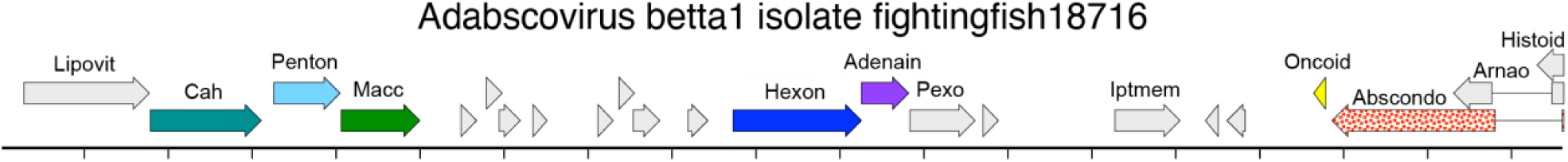
An <u>adabscovirus</u> detected in datasets representing Betta fighting fish. The inferred circular genome is linearized for display purposes. Hatch marks indicate one kilobase.

We can conceive of several hypotheses that might account for the lack of an identifiable replicase in anelloviruses and the *Betta* adabscovirus. One possibility is that they are truly replicase-less viruses that can directly recruit cellular replication factors to propagate the viral episome. Alternatively, they may encode novel replicase proteins that are currently unidentifiable or they may be multipartite viruses with a recognizable replicase encoded on an undetected genome segment. Another possibility is that they parasitize the replicases of other viruses. To address these hypotheses, anellovirus- and adabscovirus-containing datasets were intensively scanned for contigs with sequences similar to known replicases. Some anellovirus-containing datasets had short circular sequences encoding a cressvirus Rep with no apparent Cap (uCress1235, 2015, 979, 157, 1916, 1244, 1295, 1192, 1209) but other datasets only contained anellovirus sequences with no detectable replicase-like sequences. Interestingly, a herpesvirus-like sequence (<u>Herpes183851</u>) detected in adabscovirus-containing datasets has a gene we name Repoid that encodes a protein distantly similar to minichromosome maintenance superfamily 3 DNA helicases (Z=17).

Another hypothetical possibility is that seemingly replicase-less anelloviruses and adabscoviruses might encode an autocatalytic ribozyme that serves the same function as the protein-based nicking endonuclease domains that prime the rolling circle replication of cressviruses. An autocatalytic DNA-based nicking ribozyme (Chandra et al., 2009) could hypothetically account for the fact that the inferred anellovirus origin of replication (Ori) is challenging to sequence using rolling circle amplification (RCA) and standard Illumina library preparation methods (Sawaswong et al., 2019). For example, <u>Anello3206</u> has a coverage depth of 1,578x but not a single read or read pair spans the inferred Ori. Similarly, a human-associated anellovirus, uAnello3856, is covered at a depth of 41,462x but only three reads span the Ori. A testable prediction of the autocatalytic ribozyme hypothesis is that RCA products for anelloviruses might exhibit strand breaks near the Ori.

### Adenoviruses, adintoviruses, and other meldviruses

We define adintoviruses as an <u>ad</u>enovirus-like group of animal-associated viruses encoding distinctive PolB proteins bearing an N-terminal OTU-like domain and Hexon protein sequences that are reciprocally recognizable using basic BLASTp search methods (Starrett et al., 2021). Adintoviruses also encode a retrovirus-like (rve) <u>int</u>egrase sequence with a distinctive C-terminal chromodomain.

Two distinct clades of adintovirus genes, Alpha and Beta, are defined by reference sequences from *<u>Mayetiola</u>* <u>barley midge</u> and *<u>Terrapene</u>* <u>box turtle</u>, respectively. In addition to the reference Alpha-Hexon/Alpha-PolB (AA) and BB clade pairings embodied by the *Mayetiola* and *Terrapene* references a subset of adintoviruses detected in the current survey show AB or BA pairings (see isolate names).

There is a substantial degree of diversity in the inferred virion proteins of adintoviruses. Some adintoviruses do not have discernible Cah-class genes, while others encode elaborate predicted Cah proteins. In a handful of cases, HHpred searches indicate remote similarity between candidate adintovirus Cah proteins and retroviral Capsid proteins (e.g., <u>Adinto18804</u>, <u>refMeld6100</u>). There also appear to be possible rough qualitative similarities between the predicted folds and myristoylation signals of retroviral Matrix proteins and some Macc-class proteins (e.g., polyomavirus VP2 minor capsid proteins).

A great majority of adintovirus genomes encode a clear Penton gene, but a few genomes have no detectable Penton (<u>Adinto18043</u>, <u>10183</u>, <u>13229</u>, <u>14547</u>) or duplicate Pentons (<u>Adinto23690</u>, <u>18043</u>). Similarly, some adintovirus sequences appear have a C-truncated Hexon (<u>Adinto12335</u>, <u>21560</u>), while others have duplicate Hexon ORFs (<u>Adinto11680</u>, <u>21560</u>).

KnotInFrame analyses suggest that some adintovirus Penton ORFs might harbor a translational “slippery” sequence near the 3’ end that could hypothetically cause a programmed −1 ribosomal frameshift that would result in production of an isoform with an alternate C-terminal domain. Some of the alternate Penton C-termini show intriguing cysteine-rich motifs. Similar patterns are sometimes observable for adenovirus Penton, polyomavirus VP1, and papillomavirus L1 genes. A similar relationship was observed for potential <u>Hex</u>on-Aden<u>ain</u> (Hexain) fusion proteins.

Adenovirus Penton proteins typically have a distinctive domain inserted into a surface loop between the D and E strands of the β-jellyroll. The fold of the inserted domain is abstractly reminiscent of a Washington, DC-area sausage known as a <u>half smoke</u>. Although AlphaFold predictions suggest that the classic adenovirus-like DE-halfsmoke organization is typical of most adintovirus Pentons, other arrangements were observed. These include classic β-jellyroll folds with no DE loop insert (<u>Adinto14876</u>, <u>14448</u>, <u>7556</u>), DE loops with a halfsmoke insert that itself has a second halfsmoke domain inserted in an apical loop (<u>Adinto2681</u>), and triple-halfsmoke inserts (<u>Adinto5546</u>). In other cases, it appears that the Penton DE loop carries a Cah-like insert (<u>Adinto16382</u>). PDB files representing predicted Penton structures are provided in File7_PDB.zip.

Adenoviruses and adintoviruses occupy a broader consortium of eukaryotic viruses that are traditionally referred to as polinton-like, denoting to the <u>Pol</u>B and <u>Int</u>egrase genes typical of the consortium (Koonin & Krupovic, 2017). The current survey extends our prior observation (Starrett et al., 2021) of genomes uniting adenovirus-class virion proteins with genomes that lack recognizable PolB and Integrase sequences, as well as genomes that conversely unite typical adintovirus-like PolB and Integrase sequences with virion protein sequences that show predicted structural similarity to the virion proteins of larger DNA viruses, such as poxviruses and megaviruses (Figure 7). Since it is not clear to us whether these types of chimeras qualify as polinton-like, we instead apply the name <u>m</u>idsize <u>e</u>ukaryotic linear <u>D</u>NA virus (meldvirus), which accommodates the concept of widespread horizontal gene transfer among DNA virus groups.

**Figure 7:**
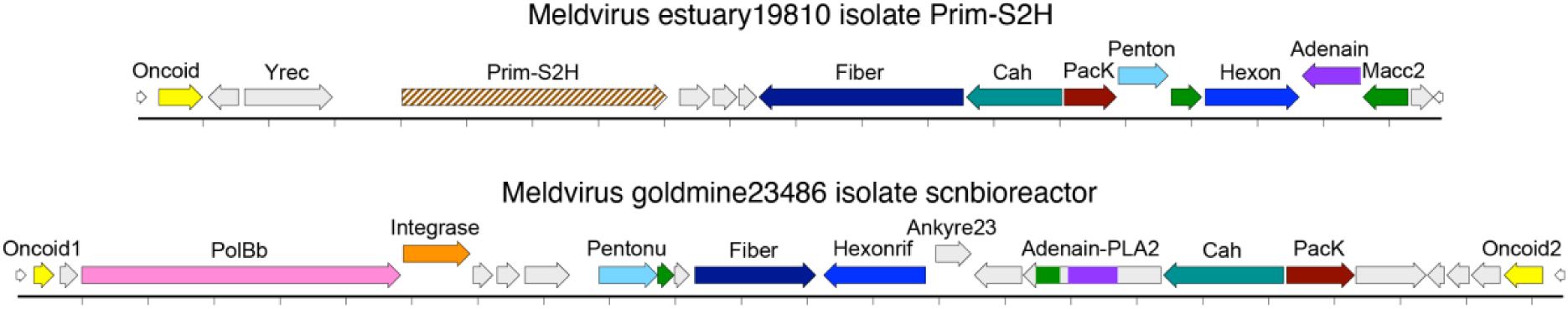
Meldvirus genomes with mixed similarities to the group traditionally referred to as polinton-like viruses. <u>Meld19810</u> unites a DNA primase-superfamily 2 helicase (Prim-S2H) and tyrosine recombinase (Yrec) with Alpha-class adintovirus virion proteins. <u>Meld23486</u> unites Beta-class adintovirus-like type B DNA polymerase (PolBb) and retrovirus-like Integrase sequences with virion protein genes that appear to be distantly related to poxvirus virion protein genes. For example, AlphaFold-predicted structures for Meld23486 Hexonrif show a Dali Z-score of 7 for <u>6BEI</u> (D13 <u>rif</u>ampicin resistance major capsid protein, a poxvirus <u>Hexon</u> homolog).

### Polyomaviruses and papillomaviruses

All currently known polyomaviruses encode their early (replicase) and late (virion protein) genes on opposing strands. A new group of polyomaviruses detected in the current survey encodes early and late genes on one strand of the circular genome. The inferred VP1 (Penton) ORF of the emerging group has a large C-terminal extension with a predicted coiled-coil structure (i.e., a candidate Cah domain). Cah-bearing polyomaviruses were detected in several spider datasets, including transcriptomic datasets representing dissected tissues (e.g., <u>Polyoma5203</u>). <u>Two examples</u> of the new polyomavirus group were detected in a <u>dataset</u> representing the NASA clean room in which the Phoenix Mars Lander was assembled (Figure 8). We speculate that the sequences might be derived from microscopic dust mites that inhabit human skin.

**Figure 8:**
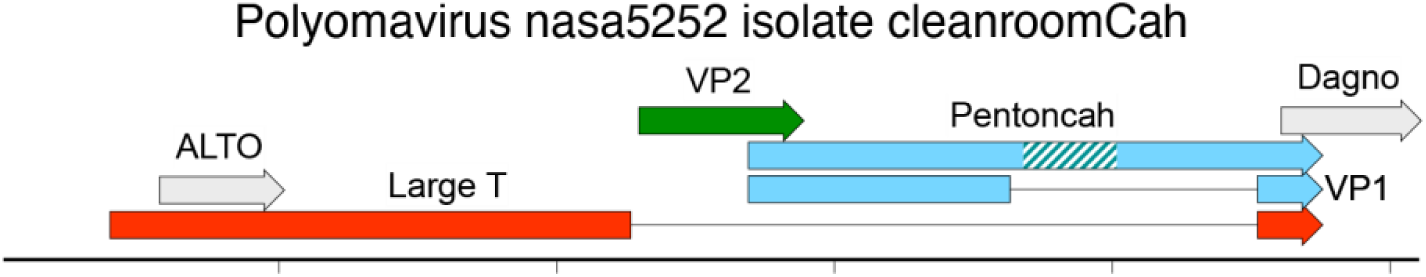
A Cah-class polyomavirus detected in a NASA clean room dataset. The hatched teal box indicates a predicted alpha helix-rich (Cah) domain.

The inferred VP2 minor capsid proteins of some of the more divergent polyomaviruses detected in the current survey (e.g., <u>Polyoma5446</u>, <u>5930</u>) show weak HHpred hits for patatin-like phospholipases (30%). This inferred <u>m</u>embrane-<u>a</u>ctive <u>c</u>apsid <u>c</u>ore (Macc) protein class is sometimes observed in the broader meldvirus consortium.

### Large DNA viruses

Although most SRA datasets were chosen for *de novo* assembly based on Diamond pre-screening for the presence of small DNA tumor virus-related sequences, the co-occurrence of herpesvirus sequences in *Betta* datasets led us to wonder whether the contig inventory might also be useful for investigating the host range of other virus types. Specifically, we hypothesized that the tumor virus screening might have inadvertently enriched for datasets representing individual animals with poor immune function, such that other virus types might also be readily detectable in the dataset.

A common human gammaherpesvirus known as Epstein-Barr virus (EBV) has been reported to productively infect dogs (Parisi et al., 2023). A high-stringency Diamond screen (E-50 cutoff) of the contig database using the sequence of EBV <u>glycoprotein B</u> as bait detected contigs with exact matches for the nucleotide sequence of EBV in dog and non-human ape datasets (see maps flagged with the term “EBV” in File2_Maps.zip). The search also revealed a sequence similar to a previously reported <u>harp seal gammaherpesvirus</u> in a polar bear <u>blood sample</u> (<u>Herpes25482</u>). The results support the view that gammaherpesviruses can sometimes be productively transmitted between distantly related host mammals.

Animal reservoirs for an emerging zoonotic pathogen known as <u>Alaskapox</u> remain uncertain. We performed Diamond screens of the contig database using Alaskapox <u>Hexon</u> as bait. Although the search did not reveal any exact matches for Alaskapox, it revealed a previously unknown chordopoxvirus (<u>Pox41634</u>) in a <u>dataset</u> representing lung material from a Cincinnati sewer rat. A lower stringency search detected an entomopoxvirus sequence (<u>Pox37252</u>) in a velvet worm <u>dataset</u>.

Elbasir and colleagues recently reported the detection of short nucleotide sequences resembling pill-bug iridescent virus 31 (<u>IIV31</u>) in datasets representing human endometrioid carcinomas (Elbasir et al., 2023). A Pubpeer <u>comment</u> argues that the IIV31-like sequences likely represent environmental contamination. To investigate this question, we performed a Diamond screen of the contig library using the IIV31 <u>Hexon</u> protein sequence as bait. IIV31-like nucleotide sequences were detected in samples of bat feces but not in any other animal datasets. An apparently complete circular genome (<u>Irido37137</u>) similar to <u>cane toad iridovirus</u> was assembled from a transcriptomic <u>dataset</u> representing the temporal polar gyrus of a 17 year-old macaque. The sequence shows no detectable nucleotide similarity to IIV31. Since the Irido37137 sequence does not show evidence of splicing, we speculate that it might represent spillover from an unknown DNA sample in the same sequencing run.

Megaviruses were originally discovered in protists (Kalafati et al., 2022). <u>Namao</u> <u>sturgeon virus</u> is the only established example of a megavirus transmitted among animals (Clouthier et al., 2020; Watson et al., 1998). The IIV31 search led to the detection of <u>Megavirus haemonchus182742</u> in a <u>dataset</u> representing barber pole worms, a common intestinal parasite of goats. Similar sequences were found in datasets representing the plant-pathogenic nematode *Globodera pallida*, in fecal samples from gorillas, humans, macaques, pigs, and rats, and in a variety of environmental samples (see files flagged with the term uMega in File2_Maps).

Mega182742 encodes two distinctive gene classes we designate <u>copi</u>es-<u>a</u>-<u>p</u>lenty (Copiap) and <u>copi</u>es <u>gal</u>ore (Copigal), which appear to have been extensively duplicated throughout the genome. Some homologs in both groups show substantial divergence, suggesting a long-term evolutionary process. A similar pattern of extensive duplications was observed for <u>leu</u>cine-rich repeat (Leurr) and <u>KilA</u>-<u>N</u>-containing (Kilan) proteins in Pox37252 and in various previously reported protist megaviruses (Machado et al., 2023). We speculate that these high-copy gene groups might be the result of an evolutionary arms race between the virus and ecdysozoan immune systems. <u>Mega</u>182742 and Pox37252 both encode group II intron-like reverse transcriptase genes (MegaRT), which could conceivably have driven the observed gene duplication patterns. Alternatively, candidate <u>F</u>anzor flap <u>endon</u>uclease homologs (Fendons) observed in Mega182742 and Irido37173 might theoretically be involved in site-specific recombination events (Saito et al., 2023).

### RNA viruses

The contig library was screened for sequences resembling the Spike and RNA-dependent RNA polymerase (RdRP) sequences of SARS-CoV-2. No sequences closely similar to SARS-CoV-2 were detected in any SRA datasets deposited prior to the year 2020.

Diamond screening using SARS-CoV-2 sequences as bait led to the detection of a divergent coronavirus-like sequence in <u>samples</u> of the gills of heat-stressed smelt fish (<u>Nidovirus hypomesus20729</u>). Interestingly, the smelt nidovirus appears to encode two distinct Spike homologs. The SARS-CoV-2 screen also detected an apparently segmented nidovirus in an *Ambystoma* salamander <u>dataset</u> (<u>Nido19192</u>). The retrieved nidovirus protein sequences were used in iterative Diamond screens of the contig library, enabling the detection of an increasingly broad range of distantly related nidoviruses in fish, amphibian, and mollusk datasets.

The iterative nidovirus <u>Spi</u>ke screen also uncovered a divergent group of chimeric <u>retro</u>virus-like sequences we refer to as spiretroviruses. It is unclear from short-read datasets whether spiretrovirus genomes are integrants with extended long terminal repeats or instead exist as circular DNA molecules. Future analyses of long-read datasets could help resolve this question.

Several chimeric RNA virus groups were detected. Tunicate and snail datasets contained tognidovirus contigs unifying a <u>tog</u>avirus-like envelope (E1) with a <u>nido</u>virus-like replicase polyprotein (<u>Tognido22085</u>, <u>20702</u>, <u>54126</u>). Contigs uniting a <u>bu</u>nyavirus-like envelope (EnvGc) with a <u>nido</u>virus-like replicase polyprotein were detected in a *Lumbricus* earthworm sample (<u>Bunido28925</u>) and a soil sample (<u>Bunido24316</u>). Another chimeric category, detected in mollusk datasets, combines a <u>nid</u>ovirus-like Spike with a t<u>oga</u>virus-like replicase polyprotein (<u>Nidoga11307</u>, <u>16890</u>). Single examples of a <u>bun</u>yavirus/<u>astro</u>virus (<u>Bunastro10698</u>) and a <u>tomb</u>usvirus/p<u>icorna</u>virus (<u>Tombicorna7041</u>) were also detected.

### Hallmark compilation

To facilitate future BLAST-based virus discovery efforts, we compiled diagnostic hallmark sequences for key virus groups examined in the current survey. Predicted proteins were clustered using an all-against-all BLASTp approach with an E-10 cutoff (File6_PolishedE10.cys). A single arbitrary exemplar from the center of each major cluster of interest was selected (File9_HallmarkProteins.fasta). An AlphaFold structure prediction for each hallmark is provided as part of File7_PDB.zip. The hallmark list is appended with concatenated core domains of hallmark small DNA tumor virus proteins.

## Discussion

Our results support the view that viruses are “the ultimate modularity” (Koonin et al., 2015). From this viewpoint, adomaviruses and papyomaviruses are not simply oddities that taxonomic systems can reasonably ignore but are instead standard examples of a basic virological principle. The evolution of highly modular organisms can be better understood using a gene-centric taxonomic approach (Figure 1).

The linear amino acid sequences of capsid protein genes tend to evolve more rapidly than the sequences of replicases (Krupovic et al., 2022). In the interest of clarity, taxonomists often choose to ignore virion proteins (Krupovic et al., 2020; Moens et al., 2017). A problem with the “Rep-only” approach is that it would lead to the misclassification of the capsid-polyphyletic cressviruses shown in Figure 2 as a single new clade of papillomaviruses. Fortunately, new artificial intelligence approaches to protein fold prediction and higher-throughput structural comparison tools have dramatically increased our power to detect distant homologs and structural/functional convergences (Mirdita et al., 2022; van Kempen et al., 2023; Villegas-Morcillo et al., 2022). In future studies, it will be important to take a more systematic approach to comparing the predicted folds of emerging capsid proteins and other hallmark viral gene classes. Experimentally determining the structures of purified virions or recombinant virion proteins will also be informative, particularly for unprecedented combinations such as the inferred tailtomavirus virion proteins. In addition to shedding light on evolutionary history, a deeper understanding of virion protein biology will open the door to the development of vaccines and antivirals.

The horizontal transfer of genes among major groups of animal viruses appears to have been rampant over million-year time scales. This is true not just for the small DNA tumor viruses we initially set out to study, but also for a group of candidate archaeal viruses carrying newly detected Hel-class DNA polymerase genes, as well as various RNA virus groups. The widespread existence of horizontal gene transfer enables a new virus-hunting approach in which known sequences are used to detect contigs that also encode previously unrecognizable viral gene classes.

We hypothesize that papillomaviruses, parvoviruses, and polyomaviruses (omaviruses) may have descended from more complex meldvirus ancestors through a series of gene-loss events. Under this hypothesis, the proposed Cah class might have served as “training wheels” during the development of the Penton-only architecture observed in familiar omaviruses. Our results suggest that some extant species may still utilize the hypothetical Cah training wheels. Another possible example of evolution through gene loss is embodied by Xeno-class adomaviruses, which appear to have replaced an ancestral Hexon with an expanded Cah. We likewise speculate that spiretroviruses might have evolved though recombination events involving meldvirus Cah, Macc, and Integrase genes and megavirus RT and Spike genes.

A more general hypothesis is that virion architecture may be more plastic than has previously been appreciated. One line of support for this idea is the remarkable range of virion morphotypes observed in electron micrographs of forest soil samples (Fischer et al., 2023). At a primary sequence level, some adintovirus species do not have detectable Cah or Penton genes, some species have highly elaborate predicted Cah or Penton genes, and some species have duplicate Cah or Penton ORFs. Similarly, some adintovirus species encode a Hexon protein that appears to be truncated after the first jellyroll domain, while others appear to encode duplicate Hexon genes. Inferred virion proteins of proposed HelPol-class maldviruses appear to have an even more dizzying degree of architectural plasticity.

Due to privacy concerns, obtaining permission to access human sequence datasets is often highly laborious. As a result, the current study examined relatively few human clinical datasets. Despite this limitation, the survey detected eight adintovirus species in human-associated datasets, including several transcriptomic datasets representing sterile tissue samples. Human-associated Beta/Beta adintoviruses are of particular interest because their PolB and Hexon protein sequences occupy a discrete vertebrate-associated clade, supporting the inference they are likely to be vertebrate-tropic, as opposed to environmental contaminants (File10_BetAdintoHexon.tre, which can be opened with <u>FigTree</u> software). We note that all adintoviruses encode a potentially mutagenic Integrase and most mammal-associated species also appear to encode homologs of anti-apoptotic genes and/or candidate oncogenes. It will be important to search for these sequences in human cancers and cancer-precursor lesions. Cancers affecting immunosuppressed individuals are of particular interest (Kooshesh et al., 2023; Starrett et al., 2023).

Understanding the virion protein biology of adintoviruses and other emerging small DNA tumor virus groups will facilitate the development of serosurveys that could address their possible prevalence. It might be possible to develop adintovirus-based gene transfer vectors (pseudoviruses) that could serve as gene delivery vehicles or as reporter systems for neutralization serology. Our observations suggest the hypothesis that hidden virion protein isoforms may be expressed via non-standard translation events that result in unexpected alternate C-terminal domains. If this hypothesis proves correct, it could be important for the development of recombinant pseudovirus systems.

An important current goal is the development of vaccines targeting commercially important fish adomaviruses. The content of such vaccines might depend on variables such as the nature of the Penton DE loop, the presence or absence of Cah or fiber genes, the presence of candidate envelope glycoproteins (e.g., <u>Adinto14876</u>)(Widen et al., 2023), or other variables that are not yet discernible through sequence-gazing alone. Some members of the polyphyletic Macc gene category might also be suitable targets for vaccines, in much the same way that papillomavirus L2 proteins (an inferred Macc class) are a target of next-generation HPV vaccines (Huber et al., 2021).

Although our contig assembly effort was relatively large-scale, it was far from comprehensive. It would be useful if public sequence repositories could offer centralized assembly snapshots of the first 50 million reads for all deep sequencing accessions. This could conceivably include <u>de-identified public versions</u> of controlled-access datasets. In the meantime, we are optimistic that the contig library generated for the current study can serve as a useful shared resource.

## Supporting information

File 1 Naming Key (Excel file)

File 2 Virus Genome Maps (GenBank text files)

File 3 All Proteins (fasta text file)

File 4 All-Protein Network E-5 (Cytoscape file)

File 5 Polished Protein Network E-5 (Cytoscape file)

File 6 Polished Protein Network E-10 (Cytoscape file)

File 7 AlphaFold-Predicted Structures (PDB files)

File 9 Hallmark Proteins (fasta text file)

File 10 Beta Adintovirus Hexon Tree (Figtree file)

## Acknowledgements

This work was funded by the NIH Intramural Research Program, with support from the Center for Cancer Research, NCI. The survey made extensive use of the computational resources of the NIH HPC <u>Biowulf</u> cluster. The authors are grateful to Alison McBride for critical evaluation of the manuscript.

